# Fexinidazole and Corallopyronin A target *Wolbachia*-infected sheath cells present in filarial nematodes

**DOI:** 10.1101/2025.01.23.634442

**Authors:** Laura Chappell, Ricardo Peguero, William R Conner, Sommer Fowler, Brandon Cooper, Kenneth Pfarr, Achim Hoerauf, Sara Lustigman, Judy Sakanari, William Sullivan

## Abstract

The discovery of the endosymbiotic bacteria *Wolbachia* as an obligate symbiont of filarial nematodes has led to antibiotic-based treatments for filarial diseases. While lab and clinical studies have yielded promising results, recent animal studies reveal that *Wolbachia* levels may rebound following treatment with suboptimal doses of the antibiotic rifampicin. Previous work showed that a likely source of the bacterial rebound in females were dense clusters of *Wolbachia* in ovarian tissue. The number, size, and density of these *Wolbachia* clusters were not diminished despite antibiotic treatment. Here we define the cellular characteristics of the *Wolbachia* clusters in *Brugia pahangi* (wBp) and identify drugs that also target them. We have evidence that the *Wolbachia* clusters originate from newly formed sheath cells adjacent to the ovarian Distal Tip Cells. The dramatically enlarged volume of an infected sheath cell is strikingly similar to endosymbiont-induced bacteriocytes found in many insect species. Ultrastructural analysis reveals that the clustered *Wolbachia* present within the sheath cells exhibit a distinct morphology and form direct connections with the oocyte membrane and possibly the cytoplasm. This includes membrane-based channels providing a connection between *Wolbachia*-infected sheath cells and oocytes. We also determined that the *Wolbachia* within the sheath cells are either quiescent or replicating at a very low rate. Screens of known antibiotics and other drugs revealed that two drugs, Fexinidazole and Corallopyronin A, significantly reduced the number of clustered *Wolbachia* located within the sheath cells.

## Introduction

The filarial nematodes *Onchocerca volvulus, Wuchereria bancrofti,* and *Brugia malayi* are human parasites that cause onchocerciasis (African river blindness) and lymphatic filariasis (elephantiasis) (Risch et al., 2024). Together these neglected diseases afflict tens of millions with many more at risk (Mathew et al., 2020). While drugs exist that effectively target the microfilariae, identifying effective drugs that kill adult worms has been problematic. This is of particular concern because adult *O. volvulus* live and remain fertile for 10 to 15 years while adult *W. bancrofti*, *B. malayi*, and *B. timori* live and remain fertile for 6 to 8 years. Thus, curing afflicted individuals requires a multiple-year drug regimen. In the late 1970s, cytological studies revealed the presence of a bacterium in filarial nematode hypodermal chords and germline tissues (Kozek, 1977; Kozek and Marroquin, 1977) (McLaren et al., 1975). Subsequent sequence analysis demonstrated that the bacteria are *Wolbachia*, an endosymbiont widely distributed among a majority of insect species and most filarial nematode species (Sironi et al., 1995) (Porter and Sullivan, 2023). In insects, *Wolbachia* is a facultative endosymbiont, but in filarial nematodes, *Wolbachia* is an obligate endosymbiont (Porter and Sullivan, 2023) maintaining a significantly smaller genome.

Loss of *Wolbachia* through antibiotic treatment results in a significant reduction in fecundity and eventual death of the adult filarial worms (Bandi et al., 1999; Hoerauf et al., 1999). This discovery led to the use of antibiotics as an effective macrofilaricidal treatment for onchocerciasis and lymphatic filariasis (Taylor et al., 2014; Wan Sulaiman et al., 2019). Clinical trials revealed that a 4 to 6 week course of antibiotics is effective at *Wolbachia* elimination and eventual killing of the adult worms (Taylor et al., 2010; Ngwewondo et al., 2021) (Hoerauf et al., 2008).

Addressing potential rebound of *Wolbachia*, a recent study demonstrated that following a one-week treatment of the filarial nematode *B. pahangi* with rifampicin *in vivo* resulted in a 95% reduction in *Wolbachia* (wBp) load; however, eight months later, *Wolbachia* titers returned to normal levels (Gunderson et al., 2020). This study also revealed the presence of dense clusters of *Wolbachia* within the ovarian tissues that were not affected despite antibiotic treatment. Significantly, the number, size, and bacterial density of these clusters remained unchanged in the antibiotic-treated worms. Given that exposure to the antibiotic had no effect on the *Wolbachia* clusters, it was postulated that the *Wolbachia* existed in privileged sites within the ovaries and that they were the likely source of the *Wolbachia* rebound months after treatment (Gunderson, et al., 2020). As antibiotic therapy is used to treat filarial diseases and becomes more widely applied, addressing the issue of recrudescence is critical.

In the present study, we provide evidence that these *Wolbachia* clusters are located within specialized ovarian cells known as sheath cells, which we herein call *Wolbachia-* infected sheath cells. Sheath cells originate at the ovarian distal tip to ultimately encompass the syncytium of developing oocytes (**Figure 1**). Ultrastructural analyses using transmission electron microscopy (TEM) revealed that these infected sheath cells possibly interact with adjacent oocytes via cytoplasmic interdigitations, and that the oocytes appear connected to the ovarian rachis by cytoskeletal bridges. Moreover, *Wolbachia* in the sheath cells are morphologically distinct from those found in the oocytes and rachis. *Wolbachia*-infected sheath cells are strikingly similar to insect bacteriocytes, which are modified host cells that provide a rich, safe environment for the occupying endosymbiont (Luan, 2024). These observations suggest a possible route by which *Wolbachia* can repopulate worm tissues that had been depleted by antibiotics, i.e. *Wolbachia* may be using these bridges to traverse the cytoplasm from the sheath cells to the surrounding oocytes and rachis.

**Figure 1.**
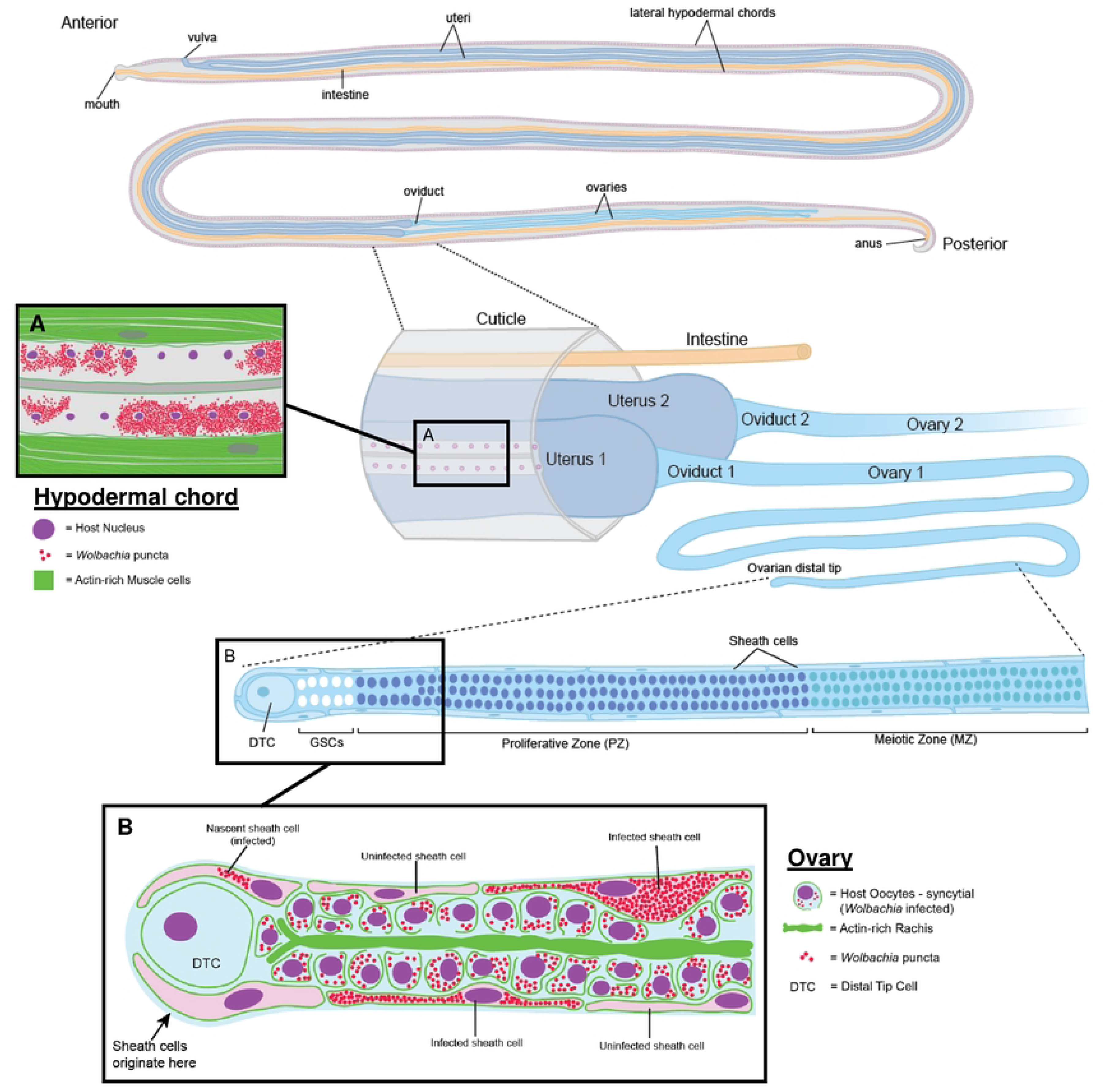
Schematic of Wolbachia-infected sheath cells in female Brugia reproductive structures. Two lateral hypodermal chords run parallel along the entire length of the nematode. These chords are syncytial with many nuclei sharing a common cytoplasm. Wolbachia are found in high densities in the hypodermal chords, as well as the germline of females. Brugia adult females maintain two germlines that continuously produce thousands of eggs. This process begins at the distal tip of the ovary in which a population of germline stem cells (GSC) associate with the distal tip cell (OTC), which serves as the stem cell niche. The GSCs produce a syncytium of mitotically active oocyte nuclei that are interconnected via a central actin-rich structure, known as the rachis. As these nuclei are propelled proximally, they leave the proliferative zone and its associated rachis, cease mitotic division, form individual cells, and begin meiotic differentiation (meiotic zone). The ovary is encompassed by a flat, elongated set of cells known as the sheath cells. These cells also originate from the OTC and migrate proximally to encompass the entire oocyte. In C. elegans, sheath cells maintain structural integrity of the oocyte and promote germline proliferation. In addition, these cells are contractile and may be involved in driving proximal migration of the oocytes. In this study, Wolbachia were found in distinct, very densely packed clusters in the sheath cells of female Brugia pahangi and Brugia malayi.

To identify new drugs that target *Wolbachia* clusters found in ovarian sheath cells, we tested the efficacy of a set of repurposed compounds. These studies revealed that two compounds, Fexinidazole and Corallopyronin A, are particularly potent in eliminating *Wolbachia* within the ovarian sheath cells.

## Results

### *Wolbachia* form dense clusters within ovarian sheath cells

Ovaries from *Brugia pahangi* female worms aged 126 to 630 days postinfection (dpi) were removed and stained with 4’,6-diamidino-2-phenylindole (DAPI), Propidium Iodide (PI), and fluorescently-labeled Phalloidin (**Figure 2**). The PI targets nucleic acids and a brief exposure preferentially stains *Wolbachia* over host cell nuclei. Unless otherwise noted, host nuclei are depicted in purple; *Wolbachia* puncta are red; and actin is green. In the examples, each cluster is localized within a host cell positioned at the outer edge of the ovary. These cells are characterized by a single flattened oblong nucleus (**Figure 2C** and **2D**, white arrowheads) and some clusters are juxtaposed (**Figure 2D**). In these infected sheath cells, the entire volume of the cytoplasm is occupied by *Wolbachia*, and the width of the cells is greatly expanded compared to uninfected cells (compare **Figure 2A, A’** to Figure **2B, B’**). In *C. elegans*, a single layer of sheath cells encompasses the germline during larval development and persists in the adult worm (Kelley and Cram, 2019). Based on the peripheral location in the gonads, the flattened shape of the nucleus, and the description by Foray et al. (2018) describing similar cells in *B. malayi*, we conclude that the *Wolbachia* clusters are localized inside sheath cells.

**Figure 2.**
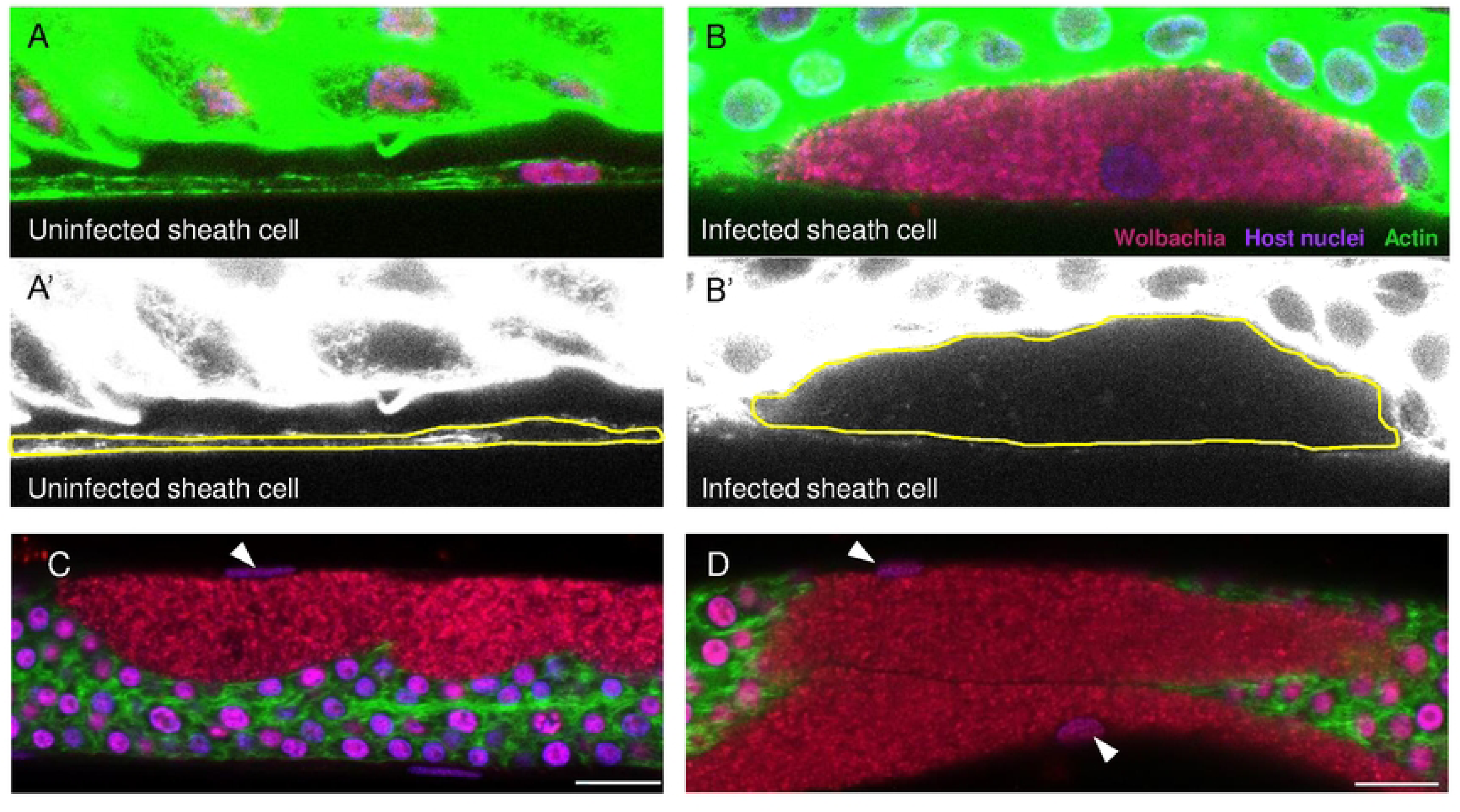
Wolbachia form dense clusters within ovarian sheath cells. **A-A’)** Confocal microscopy of a B. pahangi ovary shows a sheath cell devoid of Wolbachia. **B-B’)** A cluster of Wolbachia is seen inside a sheath cell. Yellow lines in lower panels (A’ and B’) outline the cortical actin of the sheath cells. **C-D)** Infected sheath cells are greatly expanded by Wolbachia clusters and are associated with a single, oblong nucleus (white arrowheads). All ovarian tissues are stained with Propidium Iodide (red; Wolbachia), DAPI (purple; host nuclei), and Phalloidin 488 (green; actin). All scale bars are 10 µm.

These studies were complemented by transmission electron microscopy (TEM) studies. **Figures 3A** and **3B** depict a *Wolbachia*-infected sheath cell (boxed region) closely associated with oocytes and a magnified view of the boxed region, respectively. Here the flattened sheath cell and the densely packed *Wolbachia* (black arrowheads) are evident. In the sheath cells, *Wolbachia* appear to exhibit a morphology distinct from those found in the oocytes (**Figure 3C**, black arrowhead versus black arrows). Notably, the bacteria within the sheath cell lack vacuoles, and their inner matrices appear electron lucent. While there are long regions of intact sheath cell membrane (**Figure 3B**, white arrowheads), there are also regions of disrupted membrane. Specifically, thorough ultrastructural analyses also revealed various connections between *Wolbachia*-infected sheath cells and oocytes, i.e. regions where the *Wolbachia*-infected sheath cell cytoplasm appear to have interconnections with the oocyte membrane and associated cytoplasm with apparent cytoskeletal bridges (**Figure 3B** and **3C**, asterisks, and **Figure 3E**, white arrow). Moreover, *Wolbachia* are also present in the rachis (**Figure 3E**); the rachis appears intimately associated with the surrounding oocytes including direct cytoplasmic connections between the two cell types. Confocal microscopy imaging also reveals that numerous *Wolbachia* (red dots) were present within the rachis (actin in green) as well as in the surrounding oocytes (**Figure 3F**).

**Figure 3.**
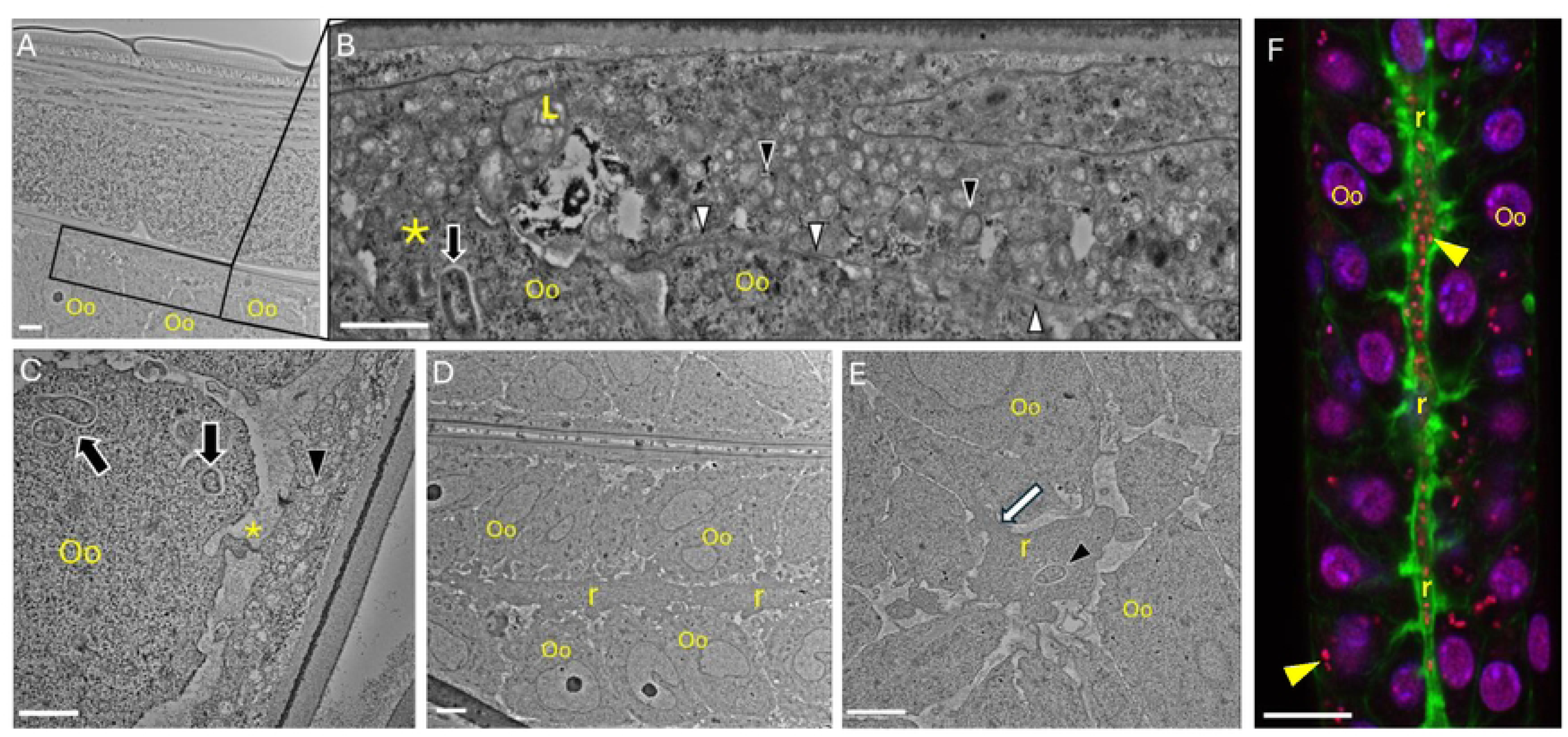
Transmission electron and confocal microscopy of adult female B. pahangi ovarian tissue. Adult female B. pahangi were cultured in vitro and processed for analysis by transmission electron and confocal microscopy. **A)** Low magnification image of ovarian tissue displaying oocytes (Oo) and an infected sheath cell (boxed region). **B)** High magnification composite image of the boxed region in panel A. An infected sheath cell containing a lysosome **(L)** and adjacent oocytes (Oo) can be seen. The sheath cell membrane (white arrowheads) is clearly visible and appears to interdigitate with an adjacent oocyte membrane (asterisk). A typical Wolbachia bacterium in a vacuole can be seen within an oocyte (black arrow). Numerous smaller, and electron lucent Wolbachia are seen within the infected sheath cell (black arrowheads). **C)** An infected sheath cell containing numerous Wolbachia (black arrowhead) is seen in proximity to an oocyte (Oo) containing several Wolbachia (black arrows). A cellular connection extending from the oocyte to the sheath cell is visible (asterisk). **D)** A low magnification image of the ovarian rachis (r) and surrounding oocytes (Oo). **E)** A high magnification image of the ovarian rachis (r) surrounded by oocytes (Oo). A bacterium (black arrowhead) can be seen in the rachis in proximity to cytoskeletal projections (white arrow) that extend from the rachis into one of the adjacent oocytes. **F)** Confocal micrograph of the ovarian rachis (r; green), surrounding oocytes (Oo; purple nuclei) and Wolbachia (yellow arrowhead; red puncta). Numerous Wolbachia can be seen throughout the rachis and within oocytes. Ovaries are stained with Propidium Iodide (red), DAPI (purple), and Phalloidin 488 (green). Scale bars: A, C-E: 2 µm; B: 1 µm; F: 10 µm.

### *Wolbachia* clusters are present in nascent sheath cells located near the ovarian Distal Tip Cell

To determine where and in which cells the *Wolbachia* clusters originate, we reasoned that cells of origin would contain smaller clusters and would not have undergone expansion. Confocal microscopy analysis reveals that cells adjacent to the Distal Tip Cell (DTC) of the ovaries occasionally contain small *Wolbachia* clusters (**Figure 4**). In *C. elegans* the DTC serves as the niche for the germline stem cells (Byrd and Kimble, 2009). Neighboring the DTC are the first of five pairs of sheath cells that divide, migrate, and elongate such that the entire *C. elegans* gonad is encompassed by sheath cells with a distinct oblong flattened shape (Kelley and Cram, 2019). We suspect that the oblong flattened cells with small *Wolbachia* clusters near the *B. pahangi* DTC are nascent sheath cells. **Figures 4A** through **4A’’** depict sequential confocal Z-planes cutting through the DTC of a *B. pahangi* ovary. In the top panel, arrows highlight two small clusters of *Wolbachia* in cells adjacent to the DTC, and the arrowheads highlight sheath cell nuclei (**Figure 4A**). **Panels 4A’** and **4A’’** show that the clusters closely associate with the host nucleus and reside within the cell. Imaging deeper into the distal tip tissue, clusters of *Wolbachia* are more apparent and can be seen extending along the length of the expanding sheath cell. **Panels 4B**, 4**B’** and **4B’’** depict three other examples of *Wolbachia* cluster-containing cells closely associated with the DTC. Given their position within the germline tissue and oblong flattened shape, we conclude they are sheath cells.

**Figure 4.**
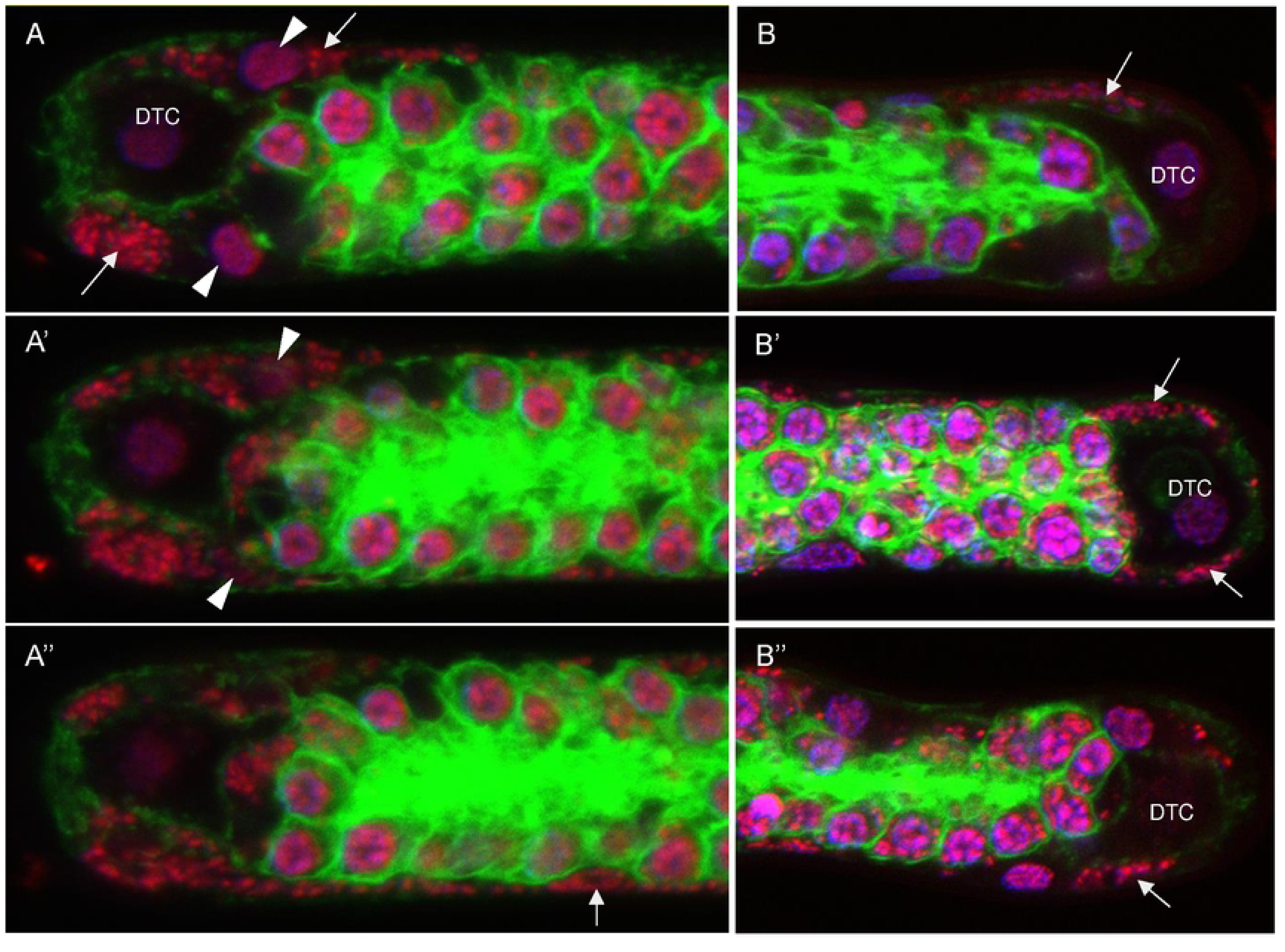
Wolbachia clusters are present in nascent sheath cells located near the ovarian Distal Tip Cell. Small Wolbachia clusters are found near, and often surrounding, the distal tip cell (OTC) of the Brugia pahangi ovary. **A-A”)** Images represent sequential Z-stacks of a nematode ovary distal tip. Two small Wolbachia clusters are seen near the OTC (A; arrows). White arrowheads point to sheath cell nuclei. Imaging deeper into the tissue reveals the bacterial cluster at the bottom edge of the ovary to be part of a larger infected sheath cell (A” arrow). Z-stack step size between images is 2.64 µm. **B-B”)** Additional images of small Wolbachia clusters found near the OTC of other B. pahangi ovaries. White arrows point to Wolbachia clusters. Tissues are stained with Propidium Iodide (red), OAPI (purple), and Phalloidin 488 (green).

### *Wolbachia*-infected sheath cells are also present in *Brugia malayi* ovarian tissues

To determine if the *Wolbachia* clusters in the sheath cells are unique to *B. pahangi,* we examined ovaries from *B. malayi*, one of three species that infects humans and causes lymphatic filariasis (Rajasekaram et al., 2017). The other two species which infect humans, *B. timori* and *W. bancrofti*, were not imaged as they cannot be developed in small animal models, and thus cultured as adult worms for laboratory experimentation. Our analyses revealed that sheath cells containing densely packed *Wolbachia* are also present in *B. malayi* ovaries (**Supplemental Figure 1**). As with *B. pahangi*, the *Wolbachia* clusters are closely associated with oblong flattened nuclei (white arrowheads) in the sheath cells that encompass the germline. Similar to *B. pahangi*, *B. malayi* clusters are also found at the edges of the ovarian tissue. Given the similarities between the two filarial species, it is likely that these *Wolbachia*-infected sheath cells are also resistant to antibiotic treatment.

### Hypodermal chord *Wolbachia,* but not oocyte or sheath cell *Wolbachia,* incorporate EdU

To determine if the *Wolbachia* are actively replicating in the mature sheath cells, we incubated ovarian tissue of adult *B. pahangi* females with 200 µM of the nucleotide analog EdU (green) for 24 and 72 hours and then fixed and stained the DNA (magenta) with a combination of PI and DAPI (**Figure 5**). Previous studies used this approach in *B. malayi* to examine DNA replication in the oocyte host nuclei (Foray et al., 2018). These studies revealed that the germline stem cells adjacent to the DTC are quiescent. However, their daughter cells, destined to form oocytes, migrate anteriorly and become mitotically active in a region known as the Proliferative Zone (PZ) as evidenced by EdU (Foray, et al., 2018).

**Figure 5.**
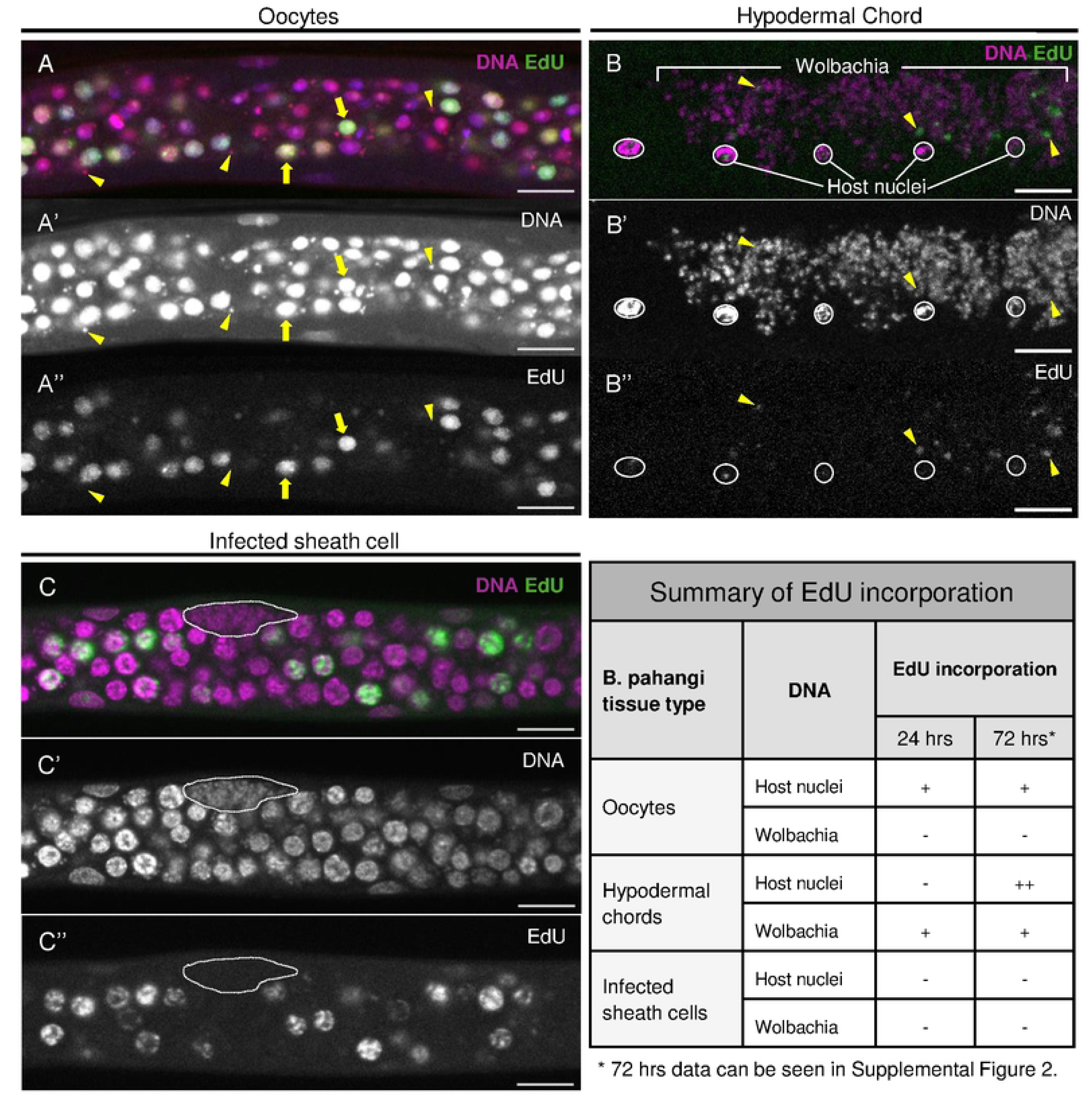
Hypodermal chord Wolbachia, but not oocyte or sheath cell Wolbachia, incorporate EdU. **A)** Ovarian tissue of adult Brugia pahangi was incubated with 200 µM EdU (green) for 24 hours. DNA is co-stained with Propidium Iodide and DAPI (magenta). Mitotically active nematode host nuclei undergoing DNA replication incorporate EdU. Yellow arrows highlight two representative nuclei that have incorporated EdU. **A’)** Single channel DNA imaging highlights Wolbachia puncta surrounding oocyte host nuclei. **A”)** Single channel EdU labeling highlights host nuclei that are undergoing or have undergone DNA replication. The three yellow arrowheads highlight Wolbachia that have not incorporated EdU. **B)** Hypodermal chords of adult Brugia pahangi were incubated with 200 µM EdU (green) for 24 hours. DNA is stained with DAPI only (magenta). Nematode host nuclei are outlined in white. All other magenta puncta are Wolbachia. EdU incorporation can be seen amongst the Wolbachia puncta. Image represents a max projection of four Z-stacks with a step size of 0.38 µm. **B’)** Single channel DNA imaging highlights the extensive distributions of Wolbachia in the hypodermal chords. **B”)** Single channel EdU labeling highlights host and bacterial DNA that has replicated or is undergoing replication. Given that EdU labels only a portion of the chromosome, it is likely the nuclei were undergoing replication during EdU incubation. C) EdU incorporation (green) was not seen in Wolbachia-infected sheath cells after 24 hours incubation at 200 µM concentration (7 total infected sheath cells were analyzed). DNA is stained with DAPI only (magenta). White dotted line indicates the outline of the infected sheath cell. **C’)** Single channel DNA imaging highlights densely packed Wolbachia clusters in the sheath cells. **C”)** Single channel EdU labeling highlights host nuclei that have replicated or are undergoing replication. Clustered Wolbachia in the sheath cells remain unlabeled. For all images, EdU is visualized with lnvitrogen Click-iT EdU imaging kit, Alexa Fluor 488. All scale bars are 10 µm.

Here we examined EdU incorporation in three host cell types: 1) the nuclei residing in the proliferative zone of the ovaries, 2) the hypodermal chord nuclei, and 3) the sheath cell nuclei (see **Figure 1**). We find that approximately a third of the oocyte Proliferative Zone host nuclei are EdU-positive (**Figure 5A**), consistent with previously published work (Foray, et al., 2018). In contrast, there was little incorporation of EdU into the hypodermal chord host nuclei after a 24hr incubation (**Figure 5B**). However, there was extensive EdU incorporation after a 72hr incubation (**Supplemental Figure 2**). No EdU incorporation was observed in host sheath cell nuclei after 24hr and 72hr incubations (**Supplemental Figure 2**).

We then examined incorporation of EdU by *Wolbachia* within the ovarian tissue, hypodermal chords, and sheath cells. We never observed EdU-positive *Wolbachia* in the Proliferative Zone (**Figure 5A**, arrowheads). Small EdU-positive puncta are observed in this zone, but these all correlate with host cell nuclei rather than *Wolbachia*. However, we did observe EdU-positive *Wolbachia* in the hypodermal chords (**Figure 5B** and **Supplemental Figure 2A,** yellow arrowheads). Nematode host nuclei, outlined in white, are EdU negative. This indicates *Wolbachia* are replicating in the hypodermal chords, in accord with previous studies (McGarry et al., 2004; Fischer et al., 2011). Finally, we examined EdU incorporation in the *Wolbachia*-infected sheath cells. Imaging the infected cells after 24hr and 72hr EdU incubations revealed that none of the clustered *Wolbachia* are EdU positive (**Figure 5C** and **Supplemental Figure 2B**). These results indicate that *Wolbachia* in the sheath cells and oocytes are not actively replicating or may be replicating at a significantly reduced rate compared to *Wolbachia* in the hypodermal chords.

### Identification of small molecules that target *Wolbachia* inside sheath cells

While a short-course (7-day) rifampicin treatment eliminated 95% of the *Wolbachia* in adult *Brugia*, this treatment had no effect on the number and size of the *Wolbachia* clusters (Gunderson, et al., 2020). In addition, rifampicin treatment did not diminish the density of *Wolbachia* in the clusters. Eight months following the 7-day course of rifampicin treatment, *Wolbachia* titers returned to normal levels. This observation led to the hypothesis that *Wolbachia* in the clusters may be a source of the rebound. Thus, we were motivated to screen for small molecules that also target *Wolbachia* within these clusters.

To this end, adult *B. pahangi* females were incubated for three days in media containing compounds at a concentration of 5 µM. Ovaries were dissected, fixed with paraformaldehyde, and stained with DAPI and PI. The dissected ovaries were then scanned using confocal microscopy and the number of *Wolbachia* clusters were counted. DMSO treated females served as a control. Because screening for drugs that eliminate the *Wolbachia* clusters using confocal microscopy is labor-intensive, we screened a limited number of compounds which consisted of a diverse set of repurposed drugs spanning a broad range of targets (**Supplemental Table 1**). As a control, we retested the effects of rifampicin, a DNA-dependent RNA polymerase inhibitor (Mosaei and Harbottle, 2019), on the frequency of the *Wolbachia* clusters.

Similar to our previous results (Gunderson, et al., 2020), while this compound effectively eliminated *Wolbachia* within the oocytes, there was no effect on the number of *Wolbachia*-infected sheath cells per ovary. The number of *Wolbachia* clusters in rifampicin-treated female worms were similar to those from the DMSO-treated controls (**Figure 6**).

**Figure 6.**
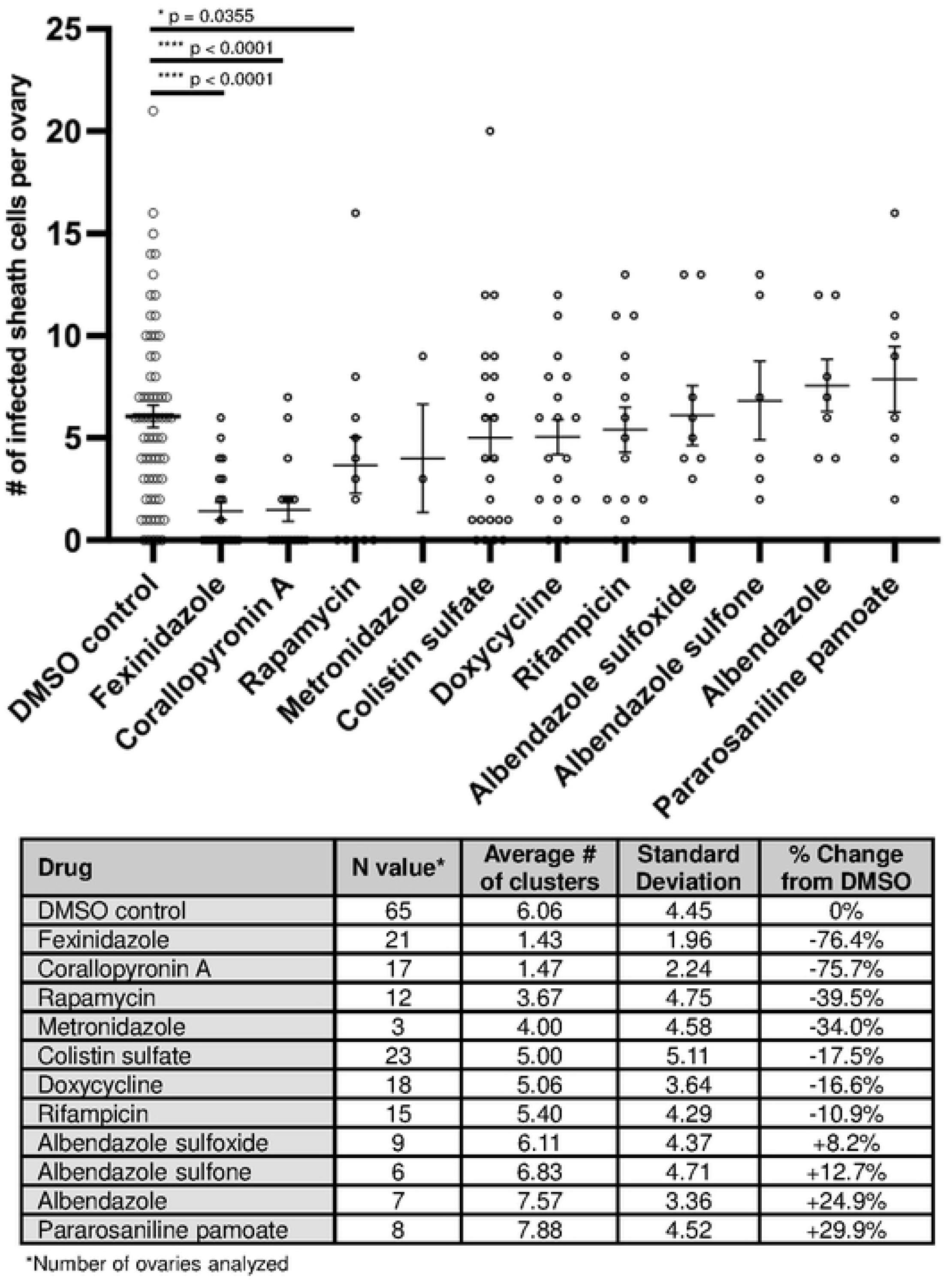
Identification of small molecules that target Wolbachia inside sheath cells. Brugia pahangi female worms were treated with various small molecule drug compounds for 72 hours at a concentration of 5 µM. Ovarian tissue from drug-treated worms was dissected, fixed and stained, and the number of Wolbachia-infected sheath cells were scored to identify the average number found per ovary. Fexinidazole, Corallopyronin A, and Rapamycin were found to significantly reduce the number of infected sheath cells in female Brugia ovaries (p < 0.05). Data points represent individual ovaries analyzed. Each ovary was stained with Pl and DAPI to visualize Wolbachia, then the entire length of the tissue was scanned from surface to surface using a confocal microscope in order to identify and score all infected sheath cells per ovary. Each compound has an experimental replicate between 2 and 5 except for Metronidazole, which was only tested once. A Mann-Whitney statistical test was used to compare each drug compound with the DMSO control (GraphPad Prism Version 10.3.1 (509)). Main horizontal lines represent means, and error bars represent standard error of the mean, SEM.

We also tested another DNA-dependent RNA polymerase inhibitor, Corallopyronin A (CorA), a proven potent anti-*Wolbachia* compound shown to be effective against a number of gram-negative and gram-positive bacteria (Schiefer et al., 2020; Krome et al., 2022). In addition, we tested two pro-drugs, fexinidazole and metronidazole (Dingsdag and Hunter, 2018; Deeks, 2019). Fexinidazole is activated by nitroreductase and the latter by oxidoreductase. Albendazole and its metabolic derivatives, Albendazole sulfone and Albendazole sulfoxide, were also tested (Borgers et al., 1975; Marriner and Bogan, 1980). Previous studies demonstrated that these compounds reduce *Wolbachia* titers in infected cell lines and *ex vivo* filarial worm studies (Serbus et al., 2012). Rapamycin is an mTOR inhibitor and potential anti-*Wolbachia* compound (Ballou and Lin, 2008; Voronin et al., 2012). Pamoate compounds were identified in a cell-based anti-*Wolbachia* screen, and Pararosaniline pamoate targets bacterial Heat Shock Protein 90 (Serbus, et al., 2012; Shahinas et al., 2015). Colistin targets Phospholipid A of the outer membrane of gram negative bacteria (Velkov et al., 2013). A summary of the compounds tested and their targets is provided in **Supplemental Table 1**.

The results of screening the above drugs are shown in **Figure 6**. Of the compounds tested, Corallopyronin A and Fexinidazole were the most potent, significantly reducing the number of *Wolbachia-*infected sheath cells per ovary by 75% and 76%, respectively, compared to the DMSO control.

## Discussion

Studies targeting *Wolbachia*, an obligate endosymbiont bacterium in human filarial parasites, have led to promising anti-*Wolbachia* based therapies for the treatment of filarial diseases. Indeed, a number of *in vivo* animal and clinical studies of infected humans demonstrated that antibiotic treatment using a long dosing regimen is an effective macrofilaricidal therapy (Fordjour and Kwarteng, 2022). However, recent *in vivo* animal studies raised concerns that the reduced titers are not always maintained after withdrawal of the anti-*Wolbachia* treatment, in particular if treatment duration is too short or dosing too low. For example, sub-optimal treatment with doxycycline (Nguyen et al., 2014) (2 weeks rather than 4 weeks) reduced *Wolbachia* in the *Litomosoides sigmodontis* infection model, but the suppression was not sustained (Hübner et al., 2019). Subsequent studies using the *B. pahangi* jird model of infection produced similar results (Gunderson, et al., 2020). In spite of a 95% reduction of *Wolbachia* following a 7-day rifampicin treatment, *Wolbachia* titers returned to normal pretreatment levels after 8 months, suggesting that *Wolbachia* may exist in a sequestered, quiescent state and may have been the source of the rebound.

Our cellular analyses of adult *B. pahangi* female worms revealed the presence of dense clusters of *Wolbachia* and that their size, number, and distribution were unaffected by exposure to rifampicin (Gunderson, et al., 2020) and doxycycline (this study). The presence of these antibiotic-resistant *Wolbachia* clusters suggested that they may account for the repopulation of *Wolbachia* in their nematode hosts. Therefore, we sought to further characterize these clusters of *Wolbachia* within the sheath cells and identify compounds that specifically target them.

Here we provide evidence that the antibiotic-resistant *Wolbachia* clusters are formed within the ovarian sheath cells of *Brugia pahangi*. In insects, some endosymbionts induce dramatic modifications to the cells in which they reside. These cells are known as bacteriocytes. They provide protection from the host’s immune system and a safe cellular environment for endosymbiont replication and transmission (Buchner, 1965; Alarcón et al., 2022). In cereal weevils, endosymbionts form distinct germline and somatic bacteriocytes (Vigneron et al., 2014). Bacteriocytes often increase the volume of their host cell, influence a wide array of cellular functions including metabolism, the immune response, and cell cycle, and provide a protective environment for the endosymbiont (Alarcón, et al., 2022). In filarial worms, the endosymbiotic bacteria *Wolbachia* reside in the hypodermal chords and in female gonadal tissues. Based on previous studies (Li et al., 2022; Foray et al. 2018) that describe sheath cells from *C. elegans* and *Brugia*, we believe that the clusters of *Wolbachia* present in the female ovaries are similar to insect bacteriocytes. Since they are found specifically in the flattened peripheral cells of the ovarian tissue, equivalent to *C. elegans* sheath cells (Kelley and Cram, 2019), we refer to the clusters of *Wolbachia* as *Wolbachia*-infected sheath cells. Interestingly, our TEM analyses suggest that *Wolbachia* may traverse between and within the ovarian tissues via cytoskeletal structures such as the rachis and other cellular connections, similar to the endosymbionts from insect bacteriocytes that have been shown to escape, migrate, and repopulate other tissues (Alarcón, et al., 2022). The potential role of cytoskeletal structures for this movement provides an explanation for why albendazole acts synergistically with some anti-*Wolbachia* drugs (Turner et al., 2017).

One of the most striking aspects of the clusters is that the load of *Wolbachia* is so extensive that the volume of the sheath cell is enlarged many-fold. In addition, *Wolbachia* inside the sheath cells appear to have a distinct morphology compared to those in the surrounding oocytes and rachis. We also observed that the *Wolbachia-* infected sheath cells have interdigitations with adjacent oocytes, and that the oocytes have cytoskeletal structures between them and the ovarian rachis that may allow *Wolbachia* to move from cell to cell. Indeed, previous studies with *B. malayi* revealed clear evidence of *Wolbachia* invading the germline from neighboring somatic cells (Landmann et al., 2010). In the present study, the observed structural junctions between the *Wolbachia*-infected sheath cells, oocytes, and rachis may allow the *Wolbachia* to traverse between cell types and extracellular matrix structures. In *C. elegans*, sheath cells form an outer monolayer of cells that encompass the oocytes and each sheath cell contains a characteristic disc-shaped flattened nucleus (Hall et al., 1999). *C. elegans* contains five pairs of sheath cells that provide structure to the oocytes (McCarter et al., 1997) as well as regulate germline proliferation (McCarter et al., 1999). In our study, we found that *Brugia* also contain sheath cells similar to *C. elegans*, but that the sheath cells surrounding the germline are infected with *Wolbachia*, making the bacteria well-positioned to invade the germline via the various cellular and structural connections.

It is unclear why the *Wolbachia* clusters in filarial nematodes are resistant to standard antibiotic treatment since insect bacteriocytes are sensitive to antibiotics (Sangaré et al., 2016). Enclosure within the ovarian sheath cells may afford *Wolbachia* a privileged site much like the granuloma surrounding *Mycobacterium tuberculosis* wherein the granuloma alters the normal internal cellular environment, resulting in hypoxia and other changes (Weeratunga et al., 2024). These changes are thought to diminish the efficacy of antibiotics that target the *Mycobacterium* (Day et al., 2023). In addition, subpopulations of *Pseudomonas aeruginosa* persist in the presence of standard antibiotics in part due to reduced metabolic activity and drug efflux (Roy et al., 2024). Support for this idea comes from our EdU incorporation studies demonstrating that the sheath cell *Wolbachia* in adult female worms (>120 days) are in a quiescent state. In filarial nematodes, *Wolbachia* undergo a major expansion during the fourth-larval stage (L4) (Fischer et al., 2014), and therefore *Wolbachia* replication in the sheath cells may be occurring sometime between the L4 and adult stage.

Clinical trials in lymphatic filariasis showed that a >95% elimination of *Wolbachia* from female filarial worms was associated with adult worm sterility and death, as seen after a 6-week doxycycline treatment, whereas a reduction by <90% (through a shorter treatment course of only 3 weeks) led to a rebound and no clear anti-parasitic activity (Debrah et al., 2006; Turner et al., 2006). Therefore, thorough elimination of the obligatory symbionts is essential in eliminating transmission of the microfilarial stage of the parasite. Because *Wolbachia* were shown to rebound in *B. pahangi* following *in vivo* short rifampicin treatment, we were motivated to identify compounds that also target *Wolbachia* within the sheath cells in an effort to ensure the prevention of the reestablishment of bacteria following the treatment. Given that the basis for the resistance of the bacteriocytes to some known antibiotics was unknown, we screened a diverse set of drugs with a broad range of targets. Two of the eleven drugs tested, Corallopyronin A and Fexinidazole, resulted in a highly significant reduction in the number of *Wolbachia*-infected sheath cells per ovary. Corallopyronin A targets bacterial DNA-dependent RNA polymerase in both gram-negative and gram-positive bacteria (Krome, et al., 2022). This compound has proven extremely effective at targeting *Wolbachia in vivo* (Schiefer, et al., 2020) and is a preclinical candidate for anti- *Wolbachia* based treatment of filarial diseases. Thus, it may be preventing the transcription of *Wolbachia* genes necessary for its survival both within and outside of the bacteriocytes and unlike other drugs, Corallopyronin A may be much more efficient at penetrating the infected sheath cells. In addition, the clustered *Wolbachia* in the sheath cells likely create a hypoxic state and unlike rifampicin, Corallopyronin A is effective in a low oxygen environment (Shima et al., 2018). Fexinidazole is an FDA-approved drug that has been particularly effective against *Trypanosoma brucei*, a blood-borne parasite and the causative agent of sleeping sickness (Jamabo et al., 2023). Known as a prodrug, it is inactive until internalized by the trypanosome. Once internalized, the trypanosome enzyme nitroreductase modifies fexinidazole, producing toxic radicals that damage parasite DNA and protein (Wittlin and Mäser, 2021). To determine if Fexinidazole might be targeting the *Wolbachia* clusters through a similar mechanism, we analyzed *Wolbachia* genomes for the presence of nitroreductase. We found that 172 of 174 *Wolbachia* genomes — including those in the filarial nematodes *O. ochengi* (wOo_09240) (Darby et al., 2012), *B. malayi* (*w*Bm_0517) (Foster et al., 2005), and *B. pahangi* (*w*Bp_RS00935) (Lebov et al., 2020) — encode a single and putatively intact nitroreductase gene copy (**Supplemental Figure 3**). The two exceptions include the nitroreductase copies observed in *w*Wb (wWb_CCY16_00998) associated with *W. bancrofti* (Chung et al., 2017) and *w*Ov (wOv_RS04175) associated with *O. volvulus* (Cotton et al., 2016) that appear disrupted (**Supplemental Figure 3**). Analysis of existing RNA-seq data from *B. malayi* ovarian proliferative (PZ) and meiotic (MZ) zones and a body-wall fragment (BW) indicate the nitroreductase is expressed at similar levels in all three tissues (Chevignon et al., 2021) (**Supplemental Figure 3**). The PZ and MZ dissections included the surrounding somatic sheath cells in which the *Wolbachia*-infected sheath cells are located. While it is not possible to determine whether the nitroreductase gene is being expressed in these cells, these studies suggest a possible mechanism of action for the effect of Fexinidazole on this *Wolbachia* population.

Anti-*Wolbachia* based therapies are proving to be one of the most effective macrofilaricidal strategies in combatting filarial nematode-based diseases. However, as with all antibiotic-based therapies, the emergence of resistant strains is a concern. For example, in insects, wild strains of *Wolbachia* differ significantly in their resistance to rifampicin (Liu et al., 2014). As described, *Wolbachia* populations rebound in filarial nematodes following withdrawal of the antibiotic if the treatment is suboptimal (with optimal conditions defined as resulting in >99% *Wolbachia* reduction in the *L. sigmodontis* infection model), a situation ripe for the selection of resistant strains. For resistant filarial *Wolbachia* to spread, they would need to enter a microfilaria larva which would then have to be picked up by a vector and be transmitted to the next human host (attrition rate approx. 10^−6^). *Wolbachia* found in the ovarian sheath cells are a likely source of the rebound and potential new resistant strains. Thus, further exploration of Corallopyronin A, Fexinidazole, and other compounds for their ability to also target *Wolbachia*-based sheath cells is warranted.

## Materials and Methods

### Parasite Material

Live *B. pahangi* and frozen *B. malayi* worms, harvested from infected jirds (*Meriones unguiculatus*), were supplied by the NIAID/NIH Filariasis Research Reagent Resource Center (FR3, Athens, GA, USA, www.filariasiscenter.org) via the Biodefense and Emerging Infections Research Resources Repository (BEI Resources, Manassas, VA, USA). The ages of the worms ranged from 126 to 630 dpi. Upon delivery, live worms were placed in an incubator at 37°C with 5% CO_2_ to recover overnight from shipping. Drug treatments or EdU incubations were performed the following day.

### Drug Treatments

Stock concentrations of 10 mM were created for all drug solutions using DMSO solvent (Sigma-Aldrich). Drugs used and their manufacturer information can be found in the table below. Corallopyronin A is a natural product purified by liquid chromatography from extracts of *Myxococcus xanthus*, a soil bacterium, and was generously gifted to us for drug screening by Achim Hoerauf and Kenneth Pfarr, Institute for Medical Microbiology, Immunology & Parasitology (IMMIP), University Hospital Bonn, Bonn, Germany.

| Drug | Manufacturer | Catalog No. |
| --- | --- | --- |
| Albendazole | Thermo Scientific | H25925-22 |
| Albendazole sulfone | Fisher Scientific | 501454432 |
| Albendazole sulfoxide | Fisher Scientific | 501415383 |
| Colistin sulfate | Fisher Scientific | 50247446 |
| Corallopyronin A | IMMIP | N/A |
| Doxycycline hydrochloride | Fisher BioReagents | BP26531 |
| Fexinidazole | Med Chem Express | HY-13801 |
| Metronidazole | WW Grainger Inc. | 30TZ38 |
| Pararosaniline pamoate | Sigma-Aldrich | SIAL-P3750 |
| Rapamycin | Fisher Scientific | 507513705 |
| Rifampicin | TCI America | R007925G |

Stock concentrations of drugs were diluted to a working concentration of 5 µM in *Brugia* media created from 80% RPMI 1640 (Gibco), 10% FBS (Life Technologies), and 10% DMEM (Corning). All drugs were used within 12 months following dilution. Worms were treated for 72 hours in 8 mL of drug-media or control media (equivalent amounts of DMSO) in 6-well plates at 37°C and 5% CO_2_. Media was refreshed daily until the final day when worms were collected in 1.5 mL of drug-media and immediately frozen at - 80°C for later immunostaining.

### Tissue Collection

For confocal analysis of germline tissue, frozen worms were thawed at room temperature and immediately fixed in 3.2% paraformaldehyde (Electron Microscopy Sciences, 15714) in PBS for 25 minutes. Worms were rinsed twice with PBS and ovaries were dissected in PBS using microdissection tweezers. Briefly, at the posterior third of the length of each worm, the outer cuticle layer was gently broken by gripping and pulling the tweezers in opposite directions at this location. Holding the tissue anterior to the cut with one tweezer, the posterior end of the tail was pulled with the other tweezer until the cuticle was completely removed, exposing the two germlines and the intestine of the worm. Dissected germlines were collected in PBS-T (1X PBS with 0.1% Triton X-100) for subsequent immunostaining.

For electron microscopy analysis of germline tissue, live worms were immediately plunged into fixative made of 2% glutaraldehyde (VWR International, 16019) and 2.5% paraformaldehyde (Electron Microscopy Sciences, 15714) buffered with 0.1M sodium cacodylate (Electron Microscopy Sciences, 11654). Worms were fixed at room temperature for 2 hours, and dissected in fixative during this incubation time. To dissect the tissue for TEM analysis, a razor blade was used to cut several 1mm posterior sections of the worm containing ovarian distal tips with attached cuticle tissue. Dissected sections, still in fixative, were placed in 4°C until shipped on ice to collaborators at New York Blood Center, NY, USA.

### Immunostaining

Dissected germline tissue was incubated with RNAse A (10 mg/mL in PBS) overnight at room temperature. Tissue was rinsed twice with PBS-T, then incubated with 1X Phalloidin 488 (Thermo Fisher) in PBS-T overnight at room temperature, protected from light. Tissue was rinsed twice with PBS-T, then stained with Propidium Iodide (Thermo Fisher) at a dilution of 1:100 in PBS-T (stock concentration of 1 mg/mL) for 25-30 seconds, and immediately rinsed twice with PBS-T. Samples were mounted on glass slides with Vectashield Mounting Media with DAPI (Vector Labs) and imaged via confocal microscopy.

### EdU Assay

5-ethynyl-2’-deoxyuridine (EdU) is a thymidine analog that incorporates into actively replicating DNA. We used the Click-iT^®^ EdU Imaging Kit with Alexa Fluor^®^ 488 from Invitrogen to perform our EdU assays. Adult female *Brugia pahangi* were incubated in 200 µM of EdU in *Brugia* media for 24 hours or 72 hours. Live worms were frozen and stored at -80°C for subsequent processing.

Frozen worms were thawed at room temperature and immediately fixed, dissected, and stained according to Invitrogen’s protocol. Briefly, worms were fixed with 3.7% formaldehyde (Electron Microscopy Sciences) in PBS for 25 minutes. Worms were rinsed twice in PBS, and germline tissue was dissected as described above in PBS. Germline tissue was collected in 0.5 mL Eppendorf tubes and washed with 3% BSA in PBS. To permeabilize membranes, tissue was incubated in 0.5% Triton X-100 in PBS for 20 minutes at room temperature. After two rinses with 3% BSA, tissue was incubated in Click-iT^®^ reaction cocktail made fresh according to protocol for 30 minutes. After two rinses with 3% BSA, tissue was mounted onto glass slides with Vectashield Mounting Media with DAPI (Vector Labs) and imaged via confocal microscopy.

### Confocal Microscopy Analysis

Images were obtained with an inverted laser scanning Leica SP5 confocal microscope using a 63x/1.4-0.6 NA oil objective and a resonant scanner (8000 Hz). For *Wolbachia*-infected sheath cell analysis, ovaries were scanned in search of *Wolbachia* clusters via direct observation through eyepieces using a DAPI filter. Ovarian issue was scanned from top surface to bottom surface along the entire length of the ovary. Due to the differential staining of PI and DAPI, *Wolbachia* can be seen as red puncta through the microscope eyepiece, making identification of clustering bacteria clear, albeit time-consuming. Number of infected sheath cells per ovary were scored manually. Digital images were processed and analyzed using ImageJ 1.54d software.

### Transmission Electron Microscopy Analysis

Six *Brugia* distal tips were received in 2% glutaraldehyde/2.5% paraformaldehyde buffered with 0.1 M sodium cacodylate. Samples were washed in buffer, and post-fixed with 1% osmium tetroxide. Samples were washed in buffer again before dehydration in an increasing ethanol series that included *en bloc* uranyl acetate staining at 70% ethanol. Following dehydration, samples were desiccated with propylene oxide and infiltrated in a 1:1 propylene oxide:epoxy resin mixture. Infiltration was continued in pure epoxy resin before embedment in pure epoxy resin at 60°C for 48 hours. Polymerized blocks were trimmed with a razor blade and ultrathin sections were collected on formvar/carbon-coated 100 mesh grids using an RMC Boeckler Powertome and a Diatome diamond knife. Sections were contrasted with uranyl acetate and Reynolds’ lead citrate and imaged in a Tecnai G2 Spirit TEM equipped with an AMT camera and imaging software. Micrograph contrast and brightness was balanced using ImageJ software.

### Statistical Analysis

Data from the drug-treatment experiments were tested for normality using the Shapiro-Wilk test and determined to significantly depart from normality. Therefore, the nonparametric equivalent of the one-way ANOVA test, the Kruskal-Wallis test, was performed on the entire dataset. It was determined that at least one sample was statistically significant from the others (p = 0.0002). A subsequent Mann-Whitney test was performed comparing the DMSO control with each drug treatment, indicating three of the drugs tested produced statistically significant results (**Figure 6**). Analyses were performed using GraphPad Prism Version 10.3.1 (509).

### Nitroreductase sequence extraction

We used the nitroreductase gene present in *Trypanosoma brucei* to serve as our target sequence (Montenegro et al., 2017). We extracted the nitroreductase domain using its UniProt annotation and used tblastn to BLAST it against the *w*Oo *Wolbachia* genome that is naturally associated with *Onchocerca ochengi* (Darby et al., 2012). This approach resulted in a single hit in the *w*Oo genome, which we extracted from NCBI’s *w*Oo annotation (wOo_09240). (This was the only gene annotated as a nitroreductase in *w*Oo.) We extracted the wOo_09240 nitroreductase domain using its UniProt annotation and used tblastn to BLAST it against all *Wolbachia* sequences in NCBI available as of Oct 11, 2023. We observed exactly one hit in each of 174 genomes, 9 of these genomes being from filarial nematodes (**Supplemental Figure 3**).

### Nitroreductase expression analysis

We obtained the RNA seq data produced by Chevignon et al. (2021) that measured *w*Bm expression in three *B. malayi* tissues: the proliferative zone, the meiotic zone, and the body wall. Following the methods of Chevignon et al. (2021), we aligned reads to the *B. malayi* mitochondria and *w*Bm *Wolbachia* genome with bowtie2 v2.5.1 (Langmead and Salzberg, 2012) We used the following options: –very-sensitive-local, —no-mixed, —no-discordant. We used featureCounts from subread v1.6.2 to count hits to each transcript (Liao et al., 2014). Normalization and differential expression analysis were performed with the R package DESeq2 v1.42.0 (Love et al., 2014).

## Acknowledgments

Funding for these studies was provided as follows: National Institutes of Health (NIH) grant <u>NIGMS-1R35GM139595</u> awarded to W.S, National Institutes of Health (NIH) grant <u>NIGMS-R35GM124701</u> awarded to B.C, Bill & Melinda Gates Foundation (OPP1017584) and the University of California, San Francisco Quantitative Biosciences Institute (QBI)-Curie/PSL for Quantitative Approaches for Studying Complex Biological Phenomena Project awarded to J.A.S., intramural funding from the New York Blood Center awarded to SL, German Center for Infection Research (DZIF, www.dzif.de) grants TTU 09.822 and 09.914 awarded to A.H and K.F. AH is a member of the Excellence Cluster Immunosensation, German Research Foundation (DFG, EXC 1023). We thank Dr. Benjamin Abrams (UCSC Life Sciences Microscopy Center, RRID: SCR_021135) for his technical support and assistance with microscopy experiments. Catharina Lindley for critical reading of the manuscript. The following reagents were provided by the NIH/NIAID Filariasis Research Reagent Resource Center for distribution through BEI Resources, NIAID, NIH: Live adult female *B. pahangi* worms (NR-48903), and frozen adult female *B. malayi* worms (NR-48893).

## Declaration of Interests

The authors declare no competing interests.

## Supplemental Information

**Supplemental Figure 1.**
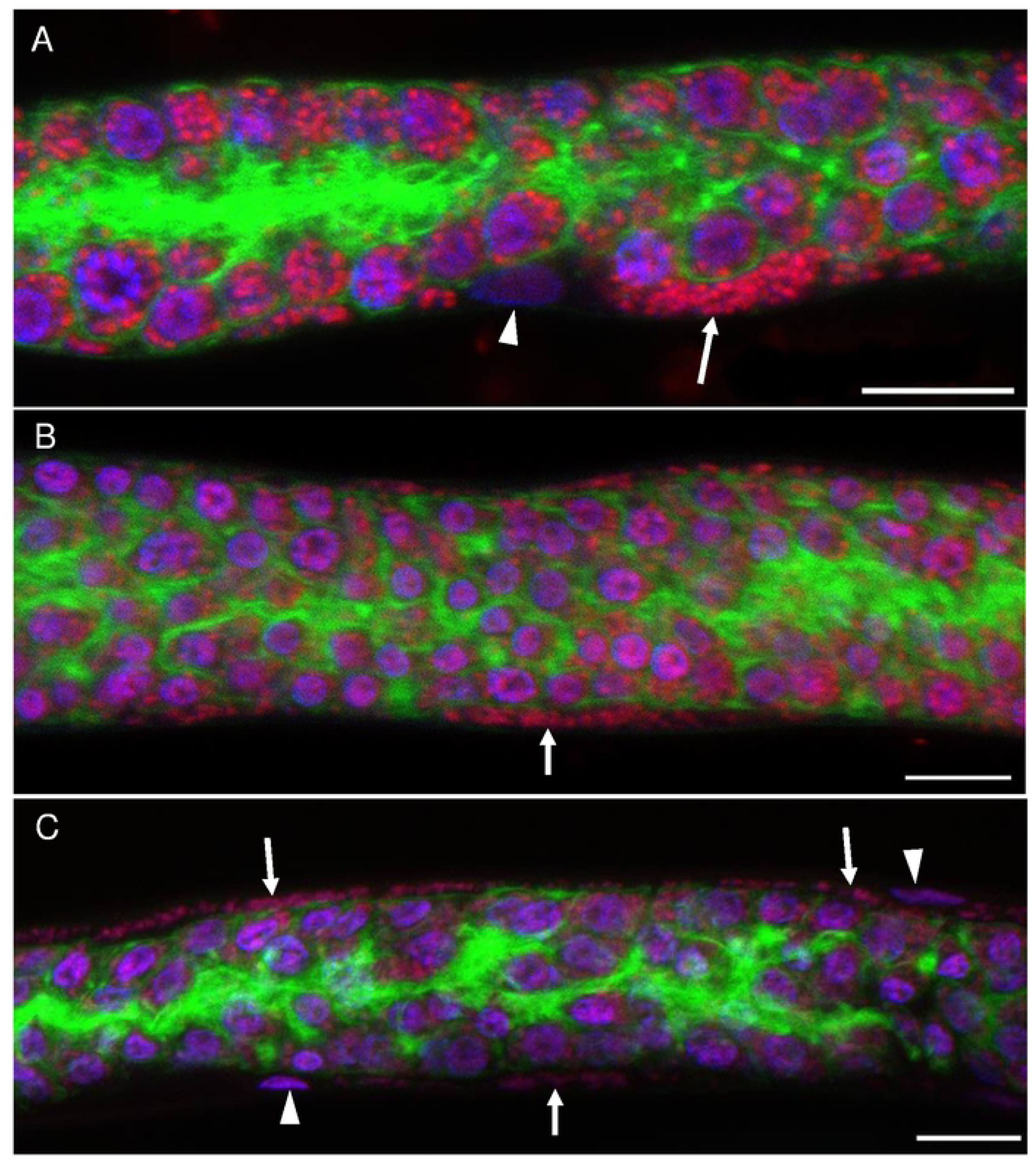
Wolbachia-infected sheath cells are also present in Brugia malayi ovarian tissues. **A-C)** Wolbachia clusters are found in one of the species of filarial nematode that infects humans, Brugia malayi. Nematode germline tissue is stained with Propidium Iodide (red), DAPI (purple), and Phalloidin 488 (green). White arrows point to Wolbachia clusters in infected sheath cells. White arrowheads point to sheath cell nuclei. All scale bars are 10µm.

**Supplemental Figure 2.**
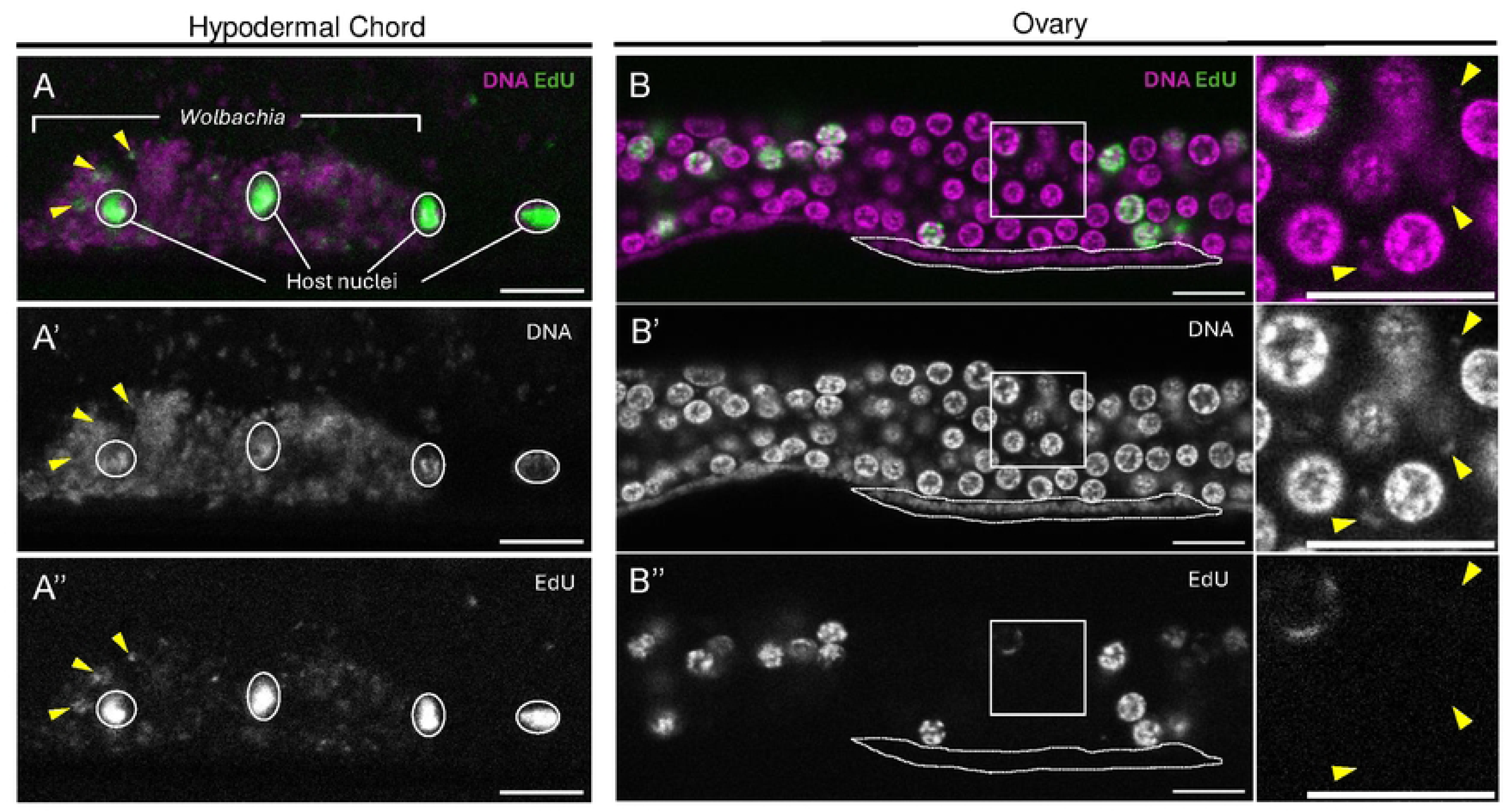
Wolbachia in the infected sheath cells do not incorporate EdU after 72 hours. **A-A”)** Hypodermal chords of adult Brugia pahangi were incubated with 200 µM EdU for 72 hours. Nematode host nuclei are outlined in white. All other magenta puncta are Wolbachia. EdU incorporation can be seen amongst the Wolbachia puncta. Image represents a max projection of four z-stacks with a step size of 0.38 µm. DNA is stained with DAPI only. **B-B”)** Ovarian tissue of adult Brugia pahangi was incubated with 200 µM EdU for 72 hours. EdU does not incorporate in Wolbachia-infected sheath cells (white dotted outline; a total of 7 infected sheath cells were analyzed). The boxed region is enlarged in the inset to the right. Nematode host oocyte nuclei incorporate EdU, but Wolbachia puncta do not (yellow arrowheads point to three representative Wolbachia puncta). DNA is stained with DAPI only. For all images, EdU is visualized with lnvitrogen Click-iT EdU imaging kit, Alexa Fluor 488. All scale bars are 10 µm.

**Supplemental Table 1.**
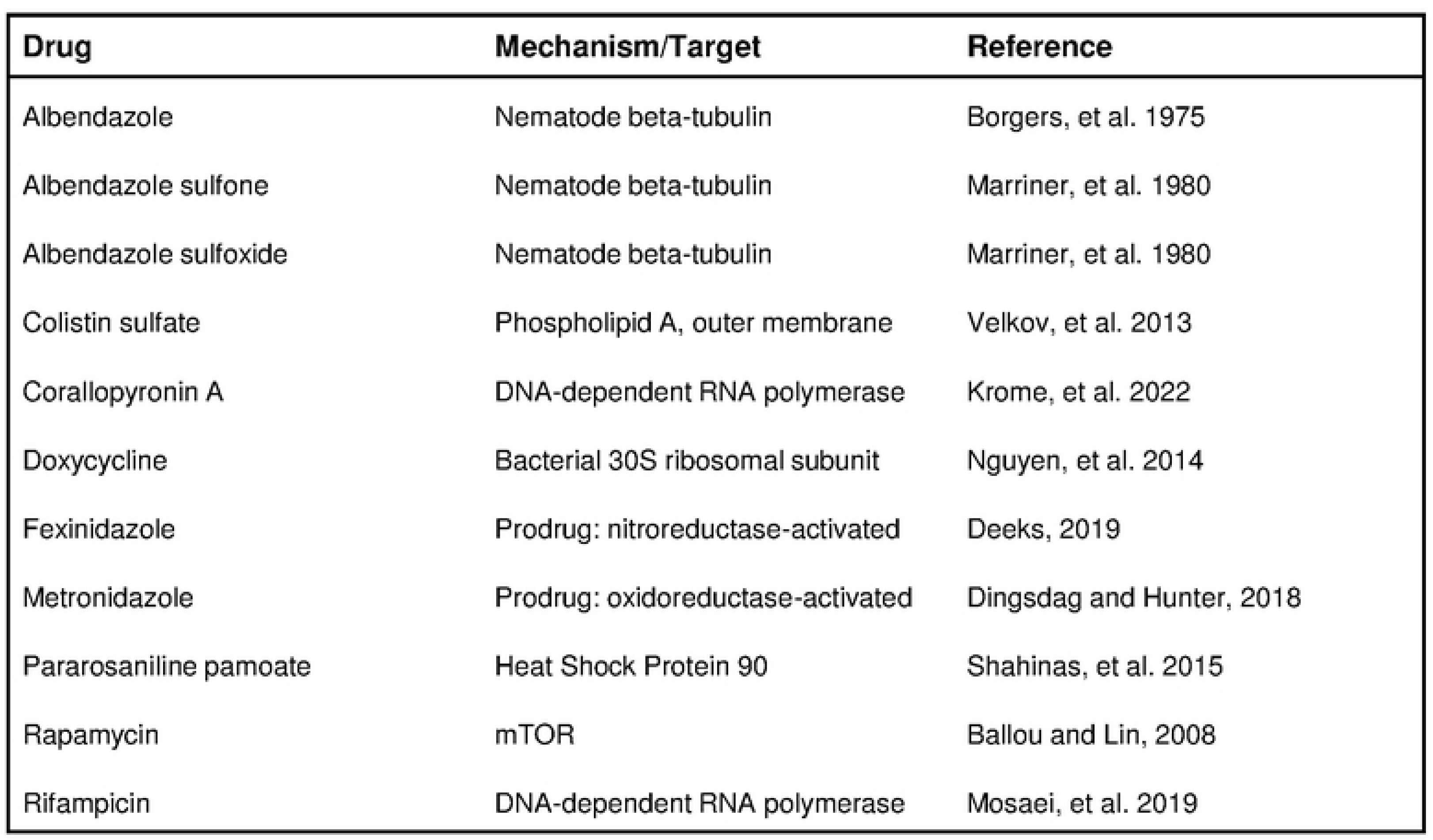
List of repurposed drugs screened for anti-wolbachial activity in infected sheath cells.

**Supplemental Figure 3.**
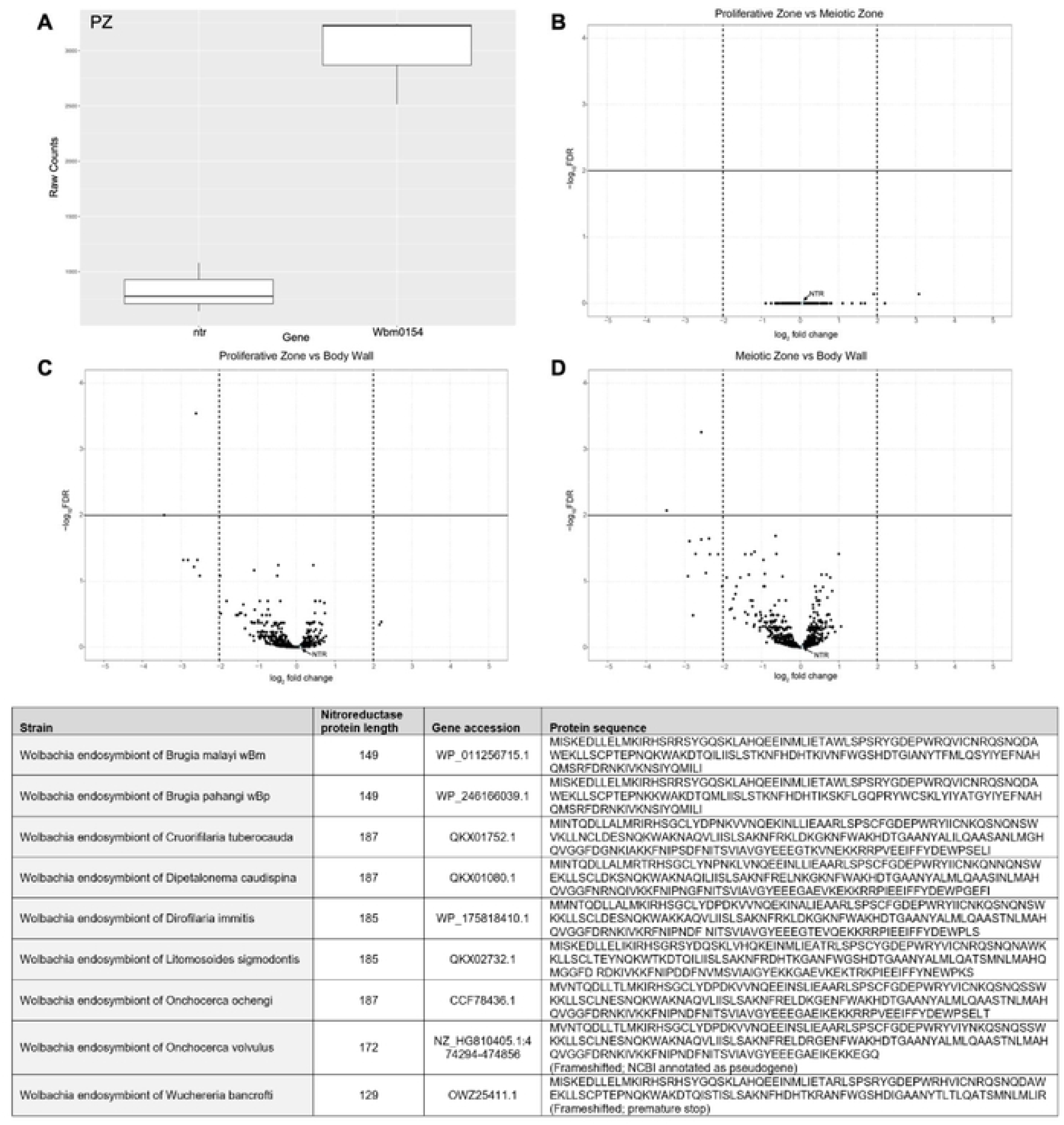
The nitroreductase gene (ntr) is expressed by Wolbachia in the nematode germline. The nitroreductase gene (ntr) is expressed at similar levels by wBm in three B. malayi host tissues: the proliferative zone (PZ), the meiotic zone (MZ), and the body wall (BW). **A)** Raw counts of ntr and actin-like gene Wbm0154 output by featureCounts for the PZ dissection. While technical aspects can affect count number, this illustrates that the ntr gene is expressed, but at a lower level relative to actin-like Wbm0154. **B-D)** Volcano plots showing similar expression of ntr across the three tissues. The Y axis is -log_10_ false discovery rate (FDR) and the X axis is log_2_ (fold change). The ntr gene is denoted in each plot. Sequencing library data were obtained from Chevignon et al. (2021). The solid black horizontal line and the vertical dashed lines denote the criteria used in Chevignon et al. (2021) for their assessment of differential gene expression: llog_2_(fold change)l>2 with an FDR <0.01. The ntr gene, like the majority in the analysis of Chevignon et al. (2021), is not differentially expressed between tissues. The relative expression of the nitroreductase is similar to expression of several other genes that include nusB (a transcription termination factor), ribosomal protein L17, and tRNA-Thr. Table indicates conserved nitroreductase genes found in the genome sequences of nine filarial nematode species.

## Notes

### Competing Interest Statement

The authors have declared no competing interest.

